# High-throughput targeted long-read single cell sequencing reveals the clonal and transcriptional landscape of lymphocytes

**DOI:** 10.1101/424945

**Authors:** Mandeep Singh, Ghamdan Al-Eryani, Shaun Carswell, James M. Ferguson, James Blackburn, Kirston Barton, Daniel Roden, Fabio Luciani, Tri Phan, Simon Junankar, Katherine Jackson, Christopher C. Goodnow, Martin A. Smith, Alexander Swarbrick

## Abstract

High-throughput single-cell RNA-Sequencing is a powerful technique for gene expression profiling of complex and heterogeneous cellular populations such as the immune system. However, these methods only provide short-read sequence from one end of a cDNA template, making them poorly suited to the investigation of gene-regulatory events such as mRNA splicing, adaptive immune responses or somatic genome evolution. To address this challenge, we have developed a method that combines targeted long-read sequencing with short-read based transcriptome profiling of barcoded single cell libraries generated by droplet-based partitioning. We use Repertoire And Gene Expression sequencing (RAGE-seq) to accurately characterize full-length T cell (TCR) and B cell (BCR) receptor sequences and transcriptional profiles of more than 7,138 lymphocytes sampled from the primary tumour and draining lymph node of a breast cancer patient. With this method we show that somatic mutation, alternate splicing and clonal evolution of T and B lymphocytes can be tracked across these tissue compartments. Our results demonstrate that RAGE-Seq is an accessible and cost-effective method for high-throughput deep single cell profiling, applicable to a wide range of biological challenges.

## Introduction

Cell diversity in humans and other vertebrates arises from complex gene rearrangement and alternative RNA splicing events that are not yet captured by current short-read sequencing technologies for measuring differential mRNA expression in single cells. A key example of this problem is the need for better ways to trace the response of single cells of the immune system during their response to cancer. Each newly-differentiated T or B lymphocyte in the immune system carries a different antigen receptor as the result of critical DNA rearrangements that alter the 450 nucleotides at the 5’ end of their T or B cell antigen receptor mRNA. In the case of B lymphocytes, they use additional DNA rearrangements to ‘isotype switch’ between 9 alternative constant region sequences comprising 1000-1500 nucleotides at the 3’ end of the heavy chain (IgH) mRNA {Di Noia, 2007 #33}, and use alternative mRNA splicing to change the 100-250 nucleotides at the 3’ end of *IGH* mRNA in order to secrete the encoded receptors as antibody {Alt, 1980 #43}. Similarly, complex gene rearrangements and alternative splicing events create pathological cell diversity amongst cancer cells. Hence there is a critical need for single cell resolution methods that capture these sequence changes occurring throughout the length of mRNA molecules, and integrate that information with gene expression features.

The extraordinary diversity of antigen receptors on B and T lymphocytes governs the development, survival and activation of these cells. T cells express on their cell surface a T cell receptor (TCR) heterodimer composed of either α and β or γ and δ chains, each the product of a different germline *TRA*, *TRB*, *TRG* or *TRD* gene loci, respectively. B cells express a B cell receptor (BCR) hetero-tetramer composed of two identical membrane immunoglobulin heavy chains encoded by the *IGH* gene locus and two identical immunoglobulin kappa or lambda light chains encoded by the *IGK* or *IGL* genes, respectively. Each of these gene loci comprise in their germline configuration a cluster of separate variable (V), diversity (D) and joining (J) gene segments, one member of each cluster becoming joined through irreversible somatic DNA rearrangements during T or B lymphocyte development in a process known as V(D)J recombination {Bassing, 2002}. Further diversity between cells is created by random addition or removal of nucleotides at the V(D)J junctions, which lie ~400 nucleotides from the 5’ end of the mRNA and encode complementarity determining region 3 (CDR3) in the antigen binding site of the receptor. The resulting diversity of the lymphocyte antigen receptor repertoire is estimated at >10^12^ different TCR or BCR proteins governed by the rule of “one cell clone - one receptor sequence” {Calis, 2014; Laydon, 2015}. Consequently, it is extremely unlikely that two cells descended from different lymphocytes will carry the same antigen receptor sequence or ‘clonotype’. As a result, when a B cell or T cell is stimulated by antigen to divide and undergo clonal expansion, the BCR or TCR sequence serves as a unique ‘clonal barcode’ and provides information on antigen specificity and cell ancestry.

Sequencing the BCR or TCR of individual lymphocytes in parallel with their transcriptome provides high-resolution insights into the adaptive immune response in a range of disease settings such as infectious disease, autoimmune disorders and cancer. A common approach to link paired antigen receptor sequences with gene expression profiles of single lymphocytes is through the use of the full-length scRNA-seq method SmartSeq2 {Picelli, 2013}, where computational methods can reconstruct paired TCRαβ sequences or paired IgH and IgL sequences from Illumina short-reads {Afik, 2017; Eltahla, 2016; Rizzetto, 2018; Stubbington, 2016; Upadhyay, 2018}. However, SmartSeq2 generally relies on plate- or well-based microfluidics and is therefore limited in the number of cells that can be processed, typically 10s to 100s. Additionally, a large number of sequencing reads are generally required to computationally reconstruct paired antigen receptors {Rizzetto, 2017}. As such, the cost per cell is relatively high ($50-$100 USD) {Ziegenhain, 2017}. Moreover, assembly of short reads makes it difficult or impossible to decipher critical alternative splicing of mRNA segments separated by more than >500 nucleotides, as occurs in *IGH* genes.

Recent technological advancements in high-throughput scRNA-seq methods allow thousands of cells to be captured and sequenced in a relatively short time frame and at a fraction of the cost {Ziegenhain, 2017}. Such methods rely on capture of polyadenylated (polyA) mRNA transcripts followed by cDNA synthesis, pooling, amplification, library construction and Illumina 3’ cDNA sequencing. The combination of fragmentation and short-read sequencing fails to sufficiently sequence the V(D)J regions of rearranged TCR and BCR transcripts, which are located in the first 500 nucleotides at the 5’ end of the transcript. Consequently, 3’-tag scRNA-seq platforms have limited application for determining clonotypic information from large numbers of lymphocytes. Variations on this approach employing 5’ cell barcodes enable the V(D)J sequences and global gene expression to be measured {Azizi, 2008}, but don’t solve the need to integrate this information with the diversity of switching and mRNA splicing involving the 3’ end of *IGH* mRNA. Recent advances in long read sequencing technologies present a potential solution to the shortcomings of short-read sequencing. Full-length cDNA reads can encompass the entire sequence of BCR and TCR transcripts, but typically suffer from higher error rates and lower sequencing depth than short read technologies {Byrne, 2017}.

Here, we describe a rapid high-throughput method to sequence full-length transcripts using targeted capture and Oxford nanopore sequencing and link this with short-read transcriptome profiling at single cell resolution. This novel method, termed <u>R</u>epertoire <u>a</u>nd <u>G</u>ene <u>E</u>xpression by <u>seq</u>uencing (RAGE-Seq), can be applied to high-throughput droplet-based scRNA-seq workflows to accurately pair gene expression profiles with targeted full-length cDNA sequences from a large number of cells. We demonstrate the power of this method by combining transcriptome profiling with full-length antigen receptor sequence characterization from thousands of human tumour-associated lymphocytes. Using *de novo* assembly of nanopore reads, complete antigen receptor sequences were recovered at high accuracy and sensitivity, including the identification of somatic mutations from immunoglobulin full-length heavy and light chains allowing for the inference of B cell clonal evolution.

## Methods

### Patient sample

Patient tissues used in this work were collected under protocol X13-0133, HREC/13/RPAH/187. HREC approval was obtained through the SLHD (Sydney Local Health District) Ethics Committee (Royal Prince Alfred Hospital zone), and site-specific approvals were obtained for all additional sites. Written consent was obtained from all patients prior to collection of tissue and clinical data stored in a de-identified manner, following pre-approved protocols. Tissue analysis was performed under protocol x14-021, LNR/14/RPAH/155.

### Single-cell suspension preparation

Following surgical resection of tumour and lymph node from patient, samples were transferred in ice cold RPMI-1640 with 50% FCS to the laboratory to be processed. Tumour was cut into approximately 1mm^3^ pieces and dissociated as per MACS human tumour dissociation kit (Miltenyi Biotec, Australia). Lymph node was similarly processed however with digestion halted at 15 minutes. After washing twice with 2% FBS in PBS, cells were resuspended in sorting buffer and passed through 70um strainers. The Jurkat T-cell line and Ramos B-cell line were cultured in RPMI-1640 medium with 10% FCS. Monocytes were flow sorted from human peripheral blood mononuclear cells (PBMCs) using a human anti-CD14 antibody. Flow cytometric sorter (BD FACS AriaIII) was used enrich for viable cells using DAPI stain, maintaining a gating threshold which omits red blood cells. Cells were centrifuged and resuspended in PBS with 2% FCS to obtain an approximate concentration of 1000 cells/ul which was counted using a haemocytometer. Samples were always handled on ice when possible and a viability of at least ~90% for all samples were confirmed by trypan blue stain prior to capture.

### Droplet based scRNAseq (10x Genomics)

Capture was performed as per 10x Chromium Single Cell 3’ (V2 chemistry) protocol with two modifications. Two extra PCR cycles were performed to allow 1:1 Full-length cDNA split, one part for Nanopore long-read sequencing and the other for short read sequencing. An lllumina NextSeq 500 was used to sequence the transcriptome library, and the yielded raw bcl file was demultiplexed and aligned (hg38 build) using CellRanger 2.0 (10x Genomics).

### Antigen-receptor capture probe design

A target enrichment library (Roche NimbleGen) was designed by first identifying gene annotations of all functional BCR and TCR genes. For each gene, genome coordinates of their corresponding exons were obtained from the GRCh38 primary assembly. In total 644 exons were targeted by the CaptureSeq array targeting ~ 128 Kb.

### Targeted capture

Following full-length cDNA amplification, 5 cycles (cell lines) or 20 cycles (primary cells) of pre-capture PCR was performed using KAPA-Hi-Fi polymerase. Next, PCR products were purified using AMPure XP beads and 500ng-1ug of amplified cDNA was used for targeted capture using the protocol previously described in {Mercer, 2014}, with minor modifications. Post-capture cDNA library size ranged from 0.6 to 2 kb.

### SmartSeq2

SmartSeq2 was performed as described by Picelli et al. {Picelli, 2014} with the following modifications: the IS PCR primer was reduced to a 50nM final concentration and the number of PCR cycles increased to 28. Sequencing was performed on the Illumina NextSeq platform. Following sequencing, reconstruction of TCR seqeunces were performed using VDJPuzzle {Eltahla, 2016}.

### scRNAseq count matrix processing

The raw gene expression matrices were normalised and scaled using Seurat (v3.4) {Satija et al. 2015}. For the cell line capture, cells that express < 250 genes or < 1000 UMIs or that contain more than 6% UMIs derived from mitochondrial genome were excluded. To reduce doublet contamination, any cells expressing > 6500 genes were discarded, additionally removing any cells that are x5 deviated from the median gene count for that cell type. For lymph node and tumour, a threshold of < 100 genes or <500 UMIs was set to allow detection of exhausted T-cells. A principle component analysis was performed on the variable genes and by using the Jackstraw method {Satija et al. 2015}, the first principle components with a P-value < 0.01 was used for dimensional reduction (40 PC). For Tumour and Lymphnode combined analysis, Seurat’s *RunCCA* was used for cross-dataset normalization to enable subsequent comparative analyses. Cell cycle scoring was performed using scRNA cell cycle gene expression scores from {Tirosh et al. 2016}.

### Identification of major clusters and differentially expressed gene analysis

The resolution set for each tSNE analysis was determinant on the strength of annotation using well known canonical marker genes and Seurat’s *FindAllMarkers* function yielding an average expression for any particular cluster >2.0-fold higher than the average expression in other sub clusters from that cell type. Seurat’s default Wlicoxon rank sum test was used for all differentially expressed gene analysis, only returning a marker with a P-value < 0.01.

### Nanopore sequencing

Hybridisation capture cDNA libraries were prepared for long read sequencing using Oxford Nanopore Technologies’ (ONT) 1D adapter ligation sequencing kit (SQK-LSK108), with the exception of one sample that used the 1D^2^ adapter ligation kit (LSK-308). The latter was base called and considered as 1D for all subsequent steps. All samples were sequenced with R9.4.1 flowcells (FLO-MIN106), with the exception of 3/6 cell line samples that were loaded onto R9.5.1 (FLO-MIN107) flowcells (including the aforementioned LSK308 sample). Base calling was performed offline on a high-performance computing cluster using ONT’s Albacore software pipeline (version 2.2.7). A list of samples, chemistries, flowcell identification numbers, and manufacturer software versions can be found in **Supplementary Table 4.**

### Demultiplexing nanopore sequencing data

Base called fastq files were pooled for each biological sample and subjected to *ad hoc* demultiplexing using a direct sequence matching strategy (i.e. 0 mismatches and indels). Cell barcode sequences (16nt) were extracted from matched short read sequencing data, as produced by 10X Genomic’s CellRanger software. Forward and reverse-complemented cell barcode sequences were then used to demultiplex the nanopore sequencing reads. The fastq headers were modified to include barcode and UMI sequences post-demultiplexing.

### De novo assembly and error correction

As highlighted in **Supplementary Figure 1**, demultiplexed reads were grouped into distinct fastq files and subjected to *de novo* assembly, then re-aligned to the resulting contigs, after which the corrected consensus was ‘polished’ with raw nanopore sequencing data. These steps were executed on a Sun Grid Engine high-performance computing cluster.

### TCR and BCR clonotype assignment

Polished fasta files containing consensus transcript contigs for each cell barcode were subjected to IgBLAST {Ye, 2013} alignment to determine V(D)J rearrangements and BLASTN alignment {Camacho, 2009} to determine the Ig or TCR constant regions exons associated with the V(D)J. For each contig, separate IgBLAST for immunoglobulin and TCR were performed using IMGT germline gene reference datasets {Lefranc, 2015}. Amino acid sequences and location of CDR3 were defined by the conserved cysteine-104 and typtophan-118 based on the IMGT numbering system {Lefranc, 2015}. IgBLAST parameters were default with the exception of returning only a single gene segment per V(D)J loci. Text-based IgBLAST output was then parsed to tab-delimited summaries, calling gene segments, framework and complementarity determining regions, mismatches, and indels relative to germline gene segments.

### Clonotype filtering

Clonotypes that were out-of-frame or that contained stop codons, termed non-productive clonotypes, were removed unless stated otherwise. BCR clonotypes containing more than 40 mutations or TCRs with more than 5 mutations in their respective V gene segments were filtered against. Analysis of somatic hypermutation of Jurkat V regions included no filtering of clonotypes based on number of mutations (**Supplementary Fig. 4**). If a cell was assigned two different TCR clonotypes with the same V and J genes but different CDR3 amino acid sequences, the TCR clonotype with the greater number of mutations in the V region was removed. If there were no differences in V region mutations the clonotype with the lowest number of reads used in assembly was removed. For BCR clonotypes only assembly read coverage was used for filtering.

### Assignment of splice isoforms

To determine the spliced constant regions exons that were associated with the V(D)J rearrangement blastn was used to align each contig against the spliced reference exons. For the *IGHC*, both the membrane and secreted versions of each constant region were included. Tabular blastn output was parsed to call constant region for each contig using the criteria of greater than 95% coverage of the spliced constant region exons and percentage identity of more than 90%. A 90% identity threshold was used because contigs used for constant region calling were not corrected for insertions or deletions.

### Integration of clonotype with scRNA-seq

Clonotypes that define groups of cells that are likely to have arisen from clonal expansion of the same progenitor B or T cell were defined either be shared gene rearrangements using the same V and J germline gene segments with identical CDR3 amino acid sequences for the T cells, and same V and J germline gene segments with 90% identical CDR3 nucleotide sequence for B cells. Clonotypes either shared the same paired chains (e.g. heavy and light chains for BCR, and alpha/beta or gamma/delta chains for the TCR) or shared TCRβ or IGH chains.

### Read subsampling

Read subsampling was performed on 200 Jurkat and 200 Ramos cells with each cell having no less than one thousand reads. The subsampling itself was performed with the sequence analysis toolkit, seqtk version 1.0-r72 (https://github.com/lh3/seqtk), using the sample command with a seed parameter of -s123. Subsampling was performed in a stepwise manner at increments of 1000, 500, 250, 100 and 50 read depths, with the resulting subsampled fastq the next input in later rounds of subsampling.

### Determining on-target Nanopore alignments

Alignment of nanopore reads to TCR and BCR genes was performed by the alignment program Minimap2 version 2.3-r536 {Li, 2018} to a custom reference fasta sequence containing TCR and BCR constant region genes, using the ‘-x map-ont’ preset. The resulting alignments were sorted and then viewed using samtools {Li, 2009 #53} version 1.7-2-gc6125d0 (with htslib 1.7-6-g6d2bfb7) and reads flagged as unmapped, not primary or supplementary were not counted as on-target.

## Results

### Targeted multiplexed full-length lymphocyte receptor sequencing

To integrate short-read and long-read mRNA sequence analysis of thousands of single cells, we designed a strategy to split full-length single-cell 3’-tag cDNA libraries prior to fragmentation for short-read sequencing, and selectively enrich BCR and TCR transcripts using targeted hybridization capture. Targeted capture was chosen over more commonly used PCR methods for repertoire analysis {Carlson, 2013; Shugay, 2014} so that retain full-length transcripts were retained. Enriched antigen-receptor molecules are subjected to long-read Nanopore sequencing to obtain both the 3’ cell-barcode and the 5’ V(D)J sequence. In parallel, short-read Illumina sequencing to profile gene expression is conducted on the remaining cDNA. By matching the cell barcodes obtained from long-read sequencing with the cell barcodes obtained from Illumina sequencing, transcriptome profiles of individual cells can then be linked with clonotype sequence (**Fig. 1**).

**Figure 1.**
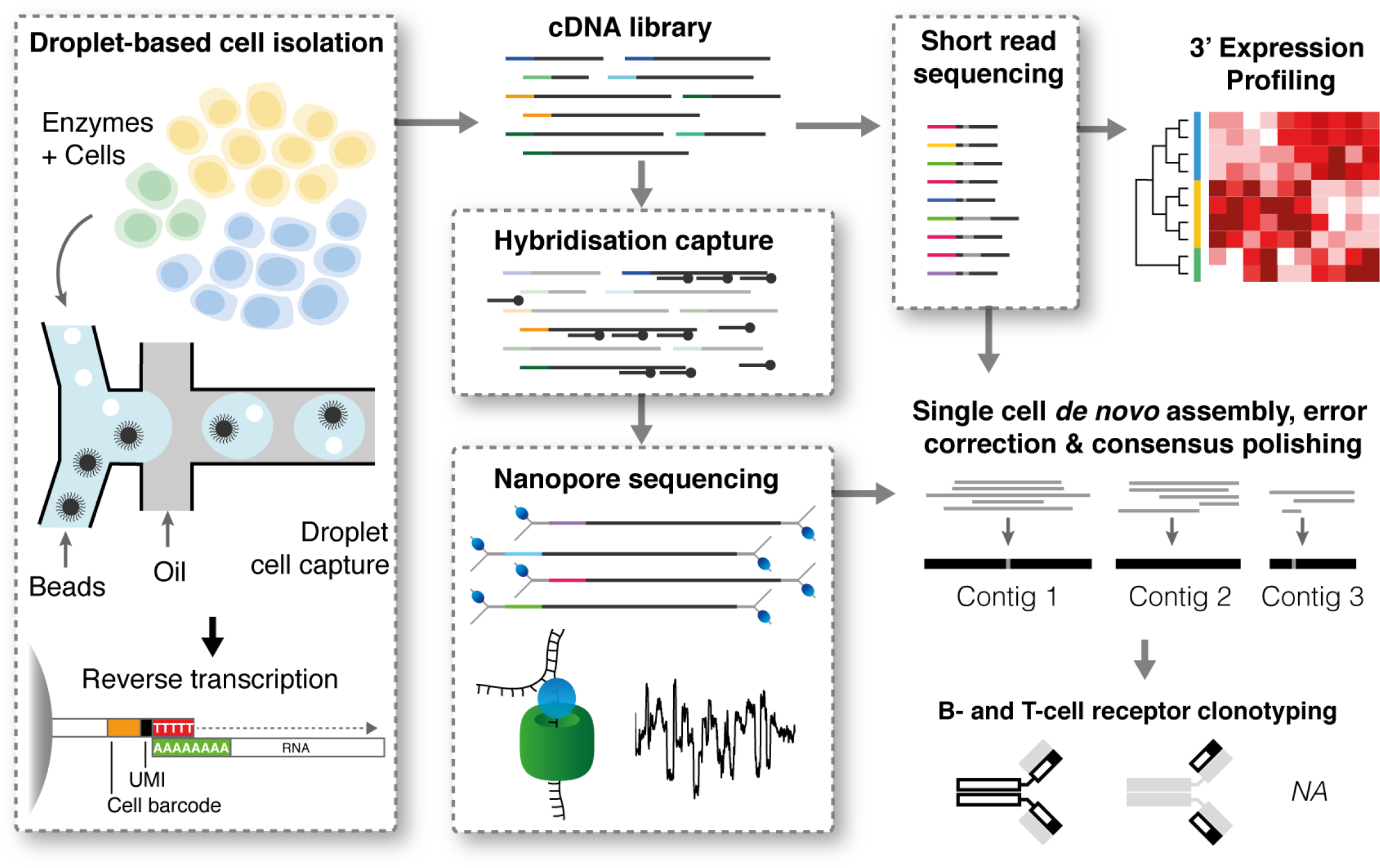
Overview of RAGE-Seq. Droplet-based single cell capture is used to generate an initial barcoded cDNA library, which is split and simultaneously subjected to (i) short read sequencing for 3’ expression profiling and (ii) hybridisation capture using custom probes followed by long read sequencing. The short read sequencing is used to cluster cellular populations and to generate high-accuracy cell barcode sequences, which are then used to demultiplex the long read data. Demultiplexed long reads are then subjected to de novo assembly, error correction, and (in this case) clonotyping analysis, resolving the complete sequence of antigen receptors with single nucleotide accuracy.

We designed a capture bait library with probes specifically targeting all annotated and functional human V, J, and Constant (C) region exons within the genomic loci that encode all TCR and BCR chains. Whole genome assemblies generated from long-read sequencing often use *de novo* assembly followed by ‘polishing’ to achieve high accuracy over 99% {Jain, 2018}. We predicted that such approaches could also be applied to Nanopore reads generated from cDNA targeted capture and developed a computational pipeline that combines *de novo* assembly with clonotype assignment on demultiplexed nanopore data to generate full-length TCR or BCR sequences for each cell (**Supplementary Fig. 1**).

### Cross-platform sequencing validation

To assess the validity of our method we performed RAGE-Seq on a mixture of the human T cell line Jurkat and the human B cell line Ramos, for which antigen receptor sequences are published {Yoshikai, 1984; Chapman, 1996} **(Supplementary Fig. 2).** A proportion of ~15% human monocyte cells were added to serve as a negative control. The dataset consisted of 1,463 Jurkat cells, 2,000 Ramos cells and 280 monocytes (**Fig. 2A, Supplementary Fig. 2**). Following nanopore sequencing, a total of 20,346,396 nanopore reads were obtained, 42.9% of which uniquely aligned to TCR and BCR constant regions (on-target reads), representing a ~13-fold enrichment when compared to non-targeted capture Illumina data (**Supplementary Fig. 2**). 18.7% of nanopore reads contained 10X cell barcodes, which recovered all barcodes identified by short-read sequencing **(Fig. 2B)**. Barcode recovery for nanopore reads that were on-target was 99.3% and 100% for Jurkat and Ramos cells, respectively **(Supplementary Fig. 2).** A strong correlation of the abundance of T cell receptor alpha constant gene (*TRAC*) reads per cell between Oxford Nanopore and Illumina sequencing was also observed (Pearson correlation = 0.79, **Fig. 2C**).

**Figure 2.**
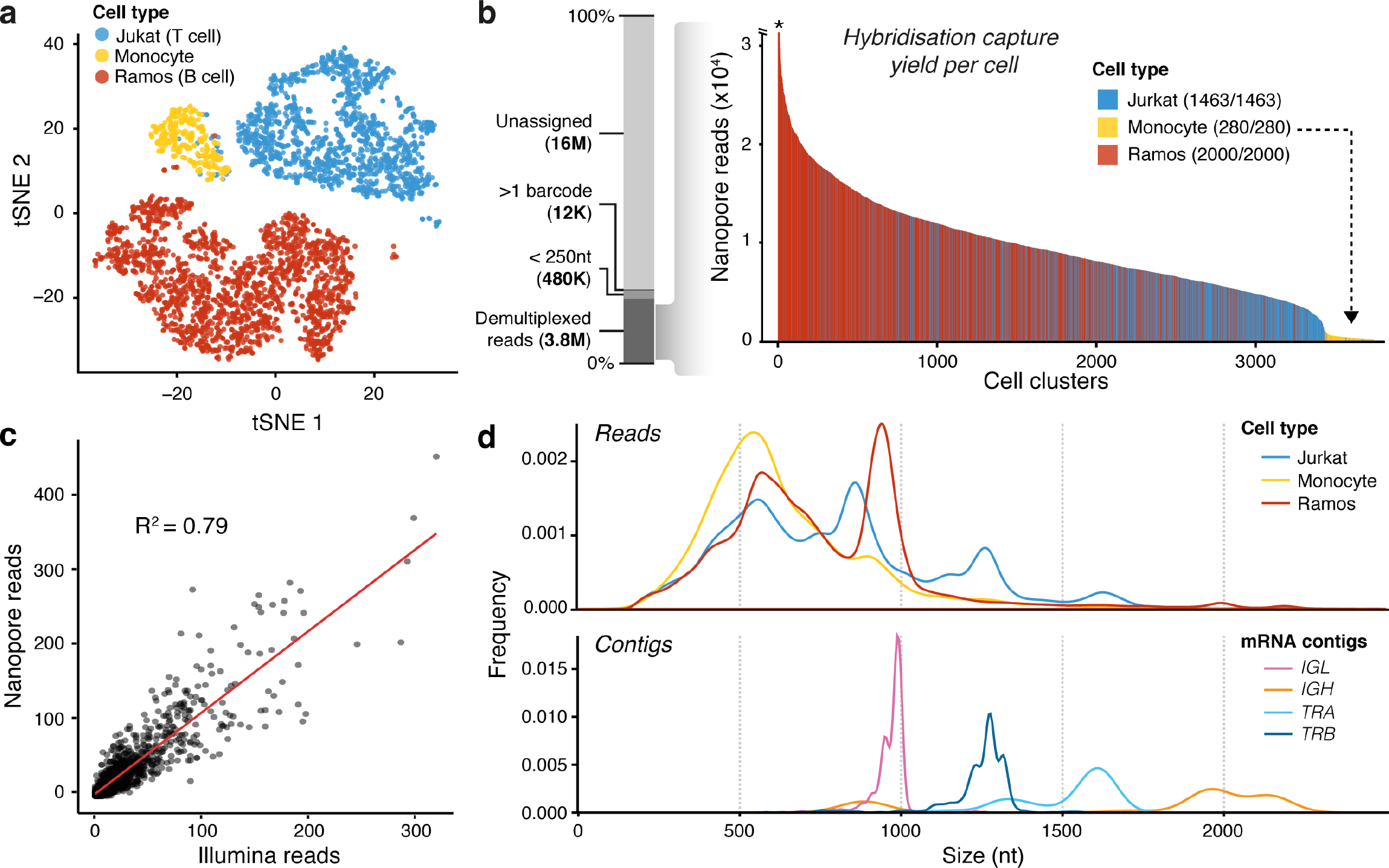
Short-read and targeted long-read single cell sequencing of immortalised B and T cell lines. (**a**) T-distributed stochastic neighbour embedding (tSNE) plot of short read sequencing data from 10X Chromium single-cell capture. Cell numbers: Jurkat=1,463; Ramos=2,000; monocytes=280 (**b**) Demultiplexing statistics for nanopore sequencing reads following targeted hybridisation capture of TCR and BCR baits. Each bar corresponds to the number of Nanopore reads per cell barcode identified with short read sequencing using exact sequence matching. Asterix indicates one cell with over 6,000 reads. Numbers next to each cell type correspond to the total number of cells with recovered barcodes. (**c**) Correlation between Illumina read counts and Oxford Nanopore read counts for T cell receptor alpha constant gene (TRAC). Each point represents an individual Jurkat cell. Pearson correlation = 0.79. (**d**) Nanopore read length distribution of demultiplexed reads assigned to cell type compared to the length distribution of polished contigs that have been assigned a productive receptor chain. Predicted nucleotide (nt) lengths of mRNA transcripts of Jurkat TRA, 1,552 nt; Jurkat TRB, 1,259 nt; Ramos IGH (secreted exons), 1,485 nt, Ramos IGH (membrane exons), 1,683 nt; Ramos IGL, 932 nt.

### Identification of receptor clonotypes

We carried out *de novo* assembly and error correction on the nanopore sequencing reads from individual cells (see Methods), generating on average 4.26 contigs per Jurkat cell, 5.24 per Ramos cell and 0.12 per Monocyte **(Supplementary Fig. 3)**. On average, 30% of contigs for Jurkat cells and 32.9% of Ramos cells were assigned a productive antigen receptor sequence. Although a minority of nanopore reads corresponded to full-length antigen receptor gene transcripts, the lengths of assembled contigs were consistent with the predicted full-length Jurkat *TRA* and *TRB* and Ramos *IGH* and *IGK* reference mRNA transcripts **(Fig. 2D, Supplementary Fig. 3).** Importantly, this shows that our *de novo* assembly approach can retain full-length transcripts.

For Jurkat cells, we recovered 18.9% (277/1,463) of cells with full-length mRNA contigs encoding paired TCRα and TCRβ chains, 13.3% with a TCRα chain only and 39.6% with a TCRβ chain only. For Ramos cells, we recovered 31% (619/2,000) of cells with nanopore contigs encoding the full coding regions for paired immunoglobulin heavy and light chains, 33% (661/2,000) with a heavy chain only and 19.1% (381/2,000) with a light chain only **(Fig. 3A)**. There was little assignment of non-reference V and J genes **(Supplementary Fig. 3).**

**Figure 3.**
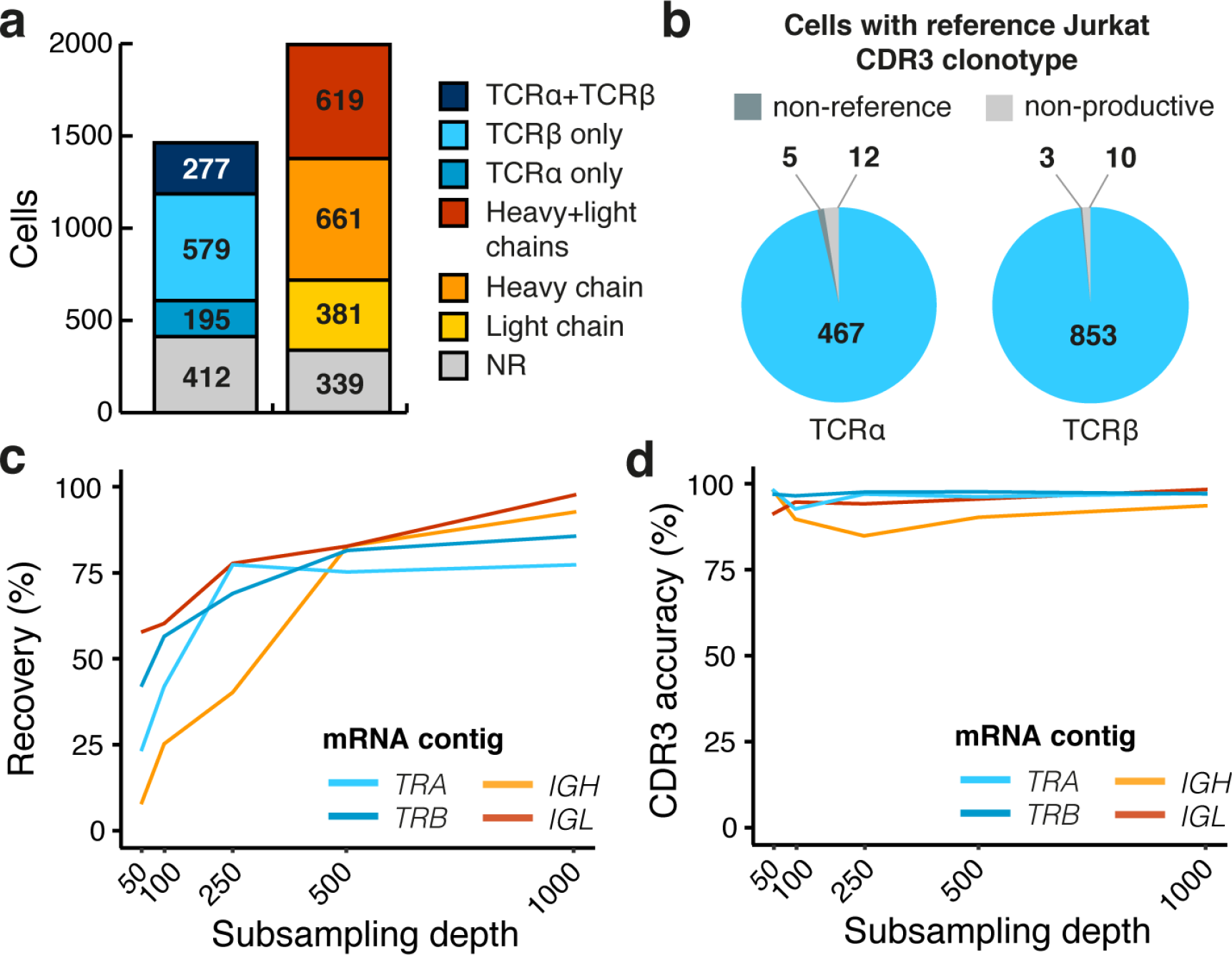
Validation of antigen receptor assembly. (**a**) Number of cells assigned productive TCR*α* and TCR *β* chains for Jurkat cells (n=1,463) or productive immunoglobulin heavy light chains for Ramos cells (n=2,000). Only receptor chains expressing the reference V and J gene combinations of Jurkat (*TRA*: *TRAV8-4*, *TRAJ3*; *TRB*: *TRBV12-3*, *TRBJ1-2*) or Ramos (*IGH*: *VH4-34*, *IGHJ6*; *IGL*: *IGLV2-14*, *IGLJ2*) were assigned. (**b**) CDR3 accuracy measured by the number of Jurkat cells with *TRA* or *TRB* sequences that directly match their reference Jurkat CDR3 nucleotide sequences (**Supplementary Fig. 2**). “On-reference” refers to a cell with a productive CDR3 sequence that does not match the reference. “Non-productive” refers to a cell with a CDR3 sequence that is out-of-frame or contains stop-codons. Only cells with reference V and J gene combinations were analysed (**Supplementary Fig. 4**). (**c**) Recovery of TCR and BCR chains as a function of sequencing depth. Subsampling was performed on 200 Jurkat and 200 Ramos cells with *>*1000 reads. For Ramos, cells with the most common *IGH* and *IGL* CDR3 sequence (**Supplementary Fig. 2**) were pre-selected. A chain is qualified as recovered if it contains its reference V and J genes. (**d**) Accuracy of the assembled CDR3 sequence versus the reference CDR3 in function of sequencing depth of sampling, as described in (c).

Next, we evaluated the accuracy of calling a correct clonotype at nucleotide resolution by investigating the CDR3 region of Jurkat cells against their known reference CDR3 sequences **(Supplementary Fig. 3)**. Of Jurkat cells with an assembled *TRA* or *TRB* contig, the percentage with the correct reference CDR3 sequence was very high. 98.9% (5/472) expressed the reference CDR3α sequence and 99.6% (853/856) expressed the reference CDR3β sequence while the number of cells carrying non-productive sequences was small (**Fig. 3B**). Assembly polishing was found to modestly increase the recovery of cells with productive chains (3.15% for TCRα and 6.14% for TCRβ) and had a small effect on the overall accuracy **(Supplementary Fig. 3)**. We also found that read depth impacted the total number of contigs recovered (**Fig. 3C**), but had little effect on the CDR3 accuracy for both TCRα and TCRβ chains and immunoglobulin heavy and light chains (**Fig. 3D**).

We also compared RAGE-Seq against the reconstruction of Jurkat TCR sequences produced using SmartSeq2 and VDJ-Puzzle {Etahla et al. 2016}. VDJ-Puzzle was able to recover a greater percentage of cells assigned TCR chains, however, RAGE-Seq proved to be ~ 30 times more cost effective on a per cell basis **(Supplementary Table 1).** Taken together, these results indicate that RAGE-Seq is both accurate and sensitive in determining clonotype sequences and has significant advantages over SmartSeq2 in terms of cost and throughput.

The Ramos cell line is known to mutate its receptors by undergoing somatic hypermutation in culture {Sale, 1998}. To identify point mutations resulting from somatic hypermutation in individual B cells, accurate sequence characterization across the entire V(D)J region of the heavy and light chain mRNA is required. Here, RAGE-Seq was able to recover 98.5% (1,259/1,278) of Ramos cells with complete *IGHV* sequences and 98.8% (988/1,000) of Ramos cells with complete *IGLV* sequences **(Supplementary Fig. 4)**. We determined amino acid replacement mutations spanning full-length V regions of the heavy and light chain from 615 Ramos cells assigned paired chains. Conserved amino acid mutations were observed in six different heavy and light chain positions, including mutations within the hydrophobic patch in framework region 1 (FR1) of the heavy chain (residues 23-25) which promotes self-reactivity {Potter, 2002} **(Fig. 4A)**. A dominant subclone within the Ramos cell line was represented by 147 cells along with 37 subclones represented by more than one cell and 319 subclones represented by a single cell **(Fig. 4A)**. We generated a clone network based on nearest neighbour distance that included the inferred germline sequence as the unmutated ancestor (**Fig. 4B**), demonstrating the evolution of individual Ramos cells undergoing active somatic hypermutation. Thus, RAGE-Seq can pair transcriptomic phenotype to immunoglobulin sequences of individual clone members within clonal populations.

**Figure 4.**
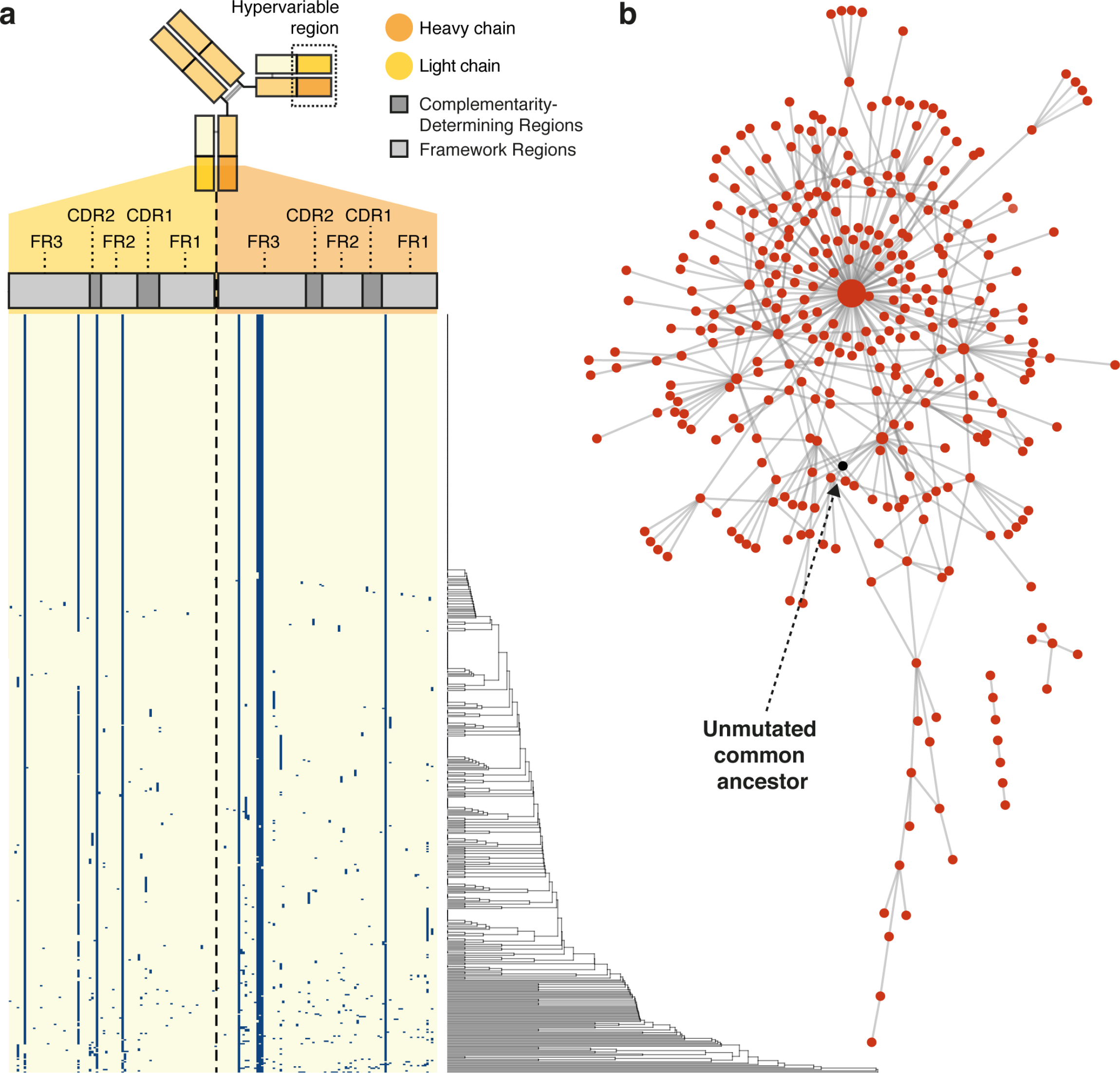
Tracking somatic hypermutation in an immortalized B-cell line. (**a**) Amino acid composition of the immunoglobulin heavy and immunoglobulin light chain V regions of Ramos cells assigned paired BCRs (n=615). Each row represents an individual cell. Blue rectangles represent non-germline amino acids, indicative of somatic hypermutation. On the right, a hierarchical clustering dendrogram of the concatenated heavy and light chain V region amino acid sequences is shown. (**b**) Network diagram of individual Ramos cells undergoing somatic hypermutation from (a), where each node corresponds to a unique heavy and light chain V(D)J sequence and the edges correspond to the number of amino acid differences between them. The largest node in the centre is the predominant sequence in this Ramos cell line represented by 147 cells, which differs from the germline reference sequence (highlighted in black, see Methods). Diagram generated with Cytoscape {Shannon et al. 2003}.

To assess the accuracy of calling point mutations we investigated Jurkat *TRAV* and *TRBV* genes, which should be completely conserved in this clonal cell line. We found only a low number of Jurkat cells with one or more inferred nucleotide mismatches to germline in these regions (*TRAV*: 5.05%, *TRBV*: 2.8%, **Supplementary Fig. 4)**.

### Analysis of lymphocytes from a human lymph node

To apply our method to primary B and T lymphocytes, we performed RAGE-Seq on a human lymph node resected from a breast cancer patient, and on a sample of the resected tumour itself. In the lymph node, we identified 4,165 T cells which could be subdivided into 6 clusters: CD4 effector memory (1,069 cells), CD4 central memory (1,321 cells), CD4 T follicular helper cells (142 cells), CD4 T regulatory cells (740 cells), CD8 central memory(487) and CD8 effector (405 cells) **(Fig. 5A, Supplementary Fig. 5)**. Among all primary T cells, we recovered 705 (16.9%) cells with paired TCRαβ chains, 1199 (28.7%) cells with a TCRα chain only and 762 (18.3%) cells with a TCRβ chain only. The recovery rate of TCR chains was comparable across the different T cell subsets **(Fig. 5B)**. We could also detect two different *TRA* or *TRB* sequences in 138 (9.5%) and 35 (1.8%) T cells, respectively, a frequency similar to previous reports {Stubbington, 2016; Eltahla, 2016}. Among the 1,619 B cells in the lymph node, we recovered 689 (42.6%) cells with paired immunoglobulin heavy and light chain contigs, 188 (11.6%) cells with only a heavy chain contig and 557 (35.6%) cells with only a light chain contig (**Fig. 5B**). Similar to the cell line experiment, all the cell barcodes were recovered across both sequencing platforms and full-length receptor chain sequences were assembled **(Supplementary Fig. 5).**

**Figure 5.**
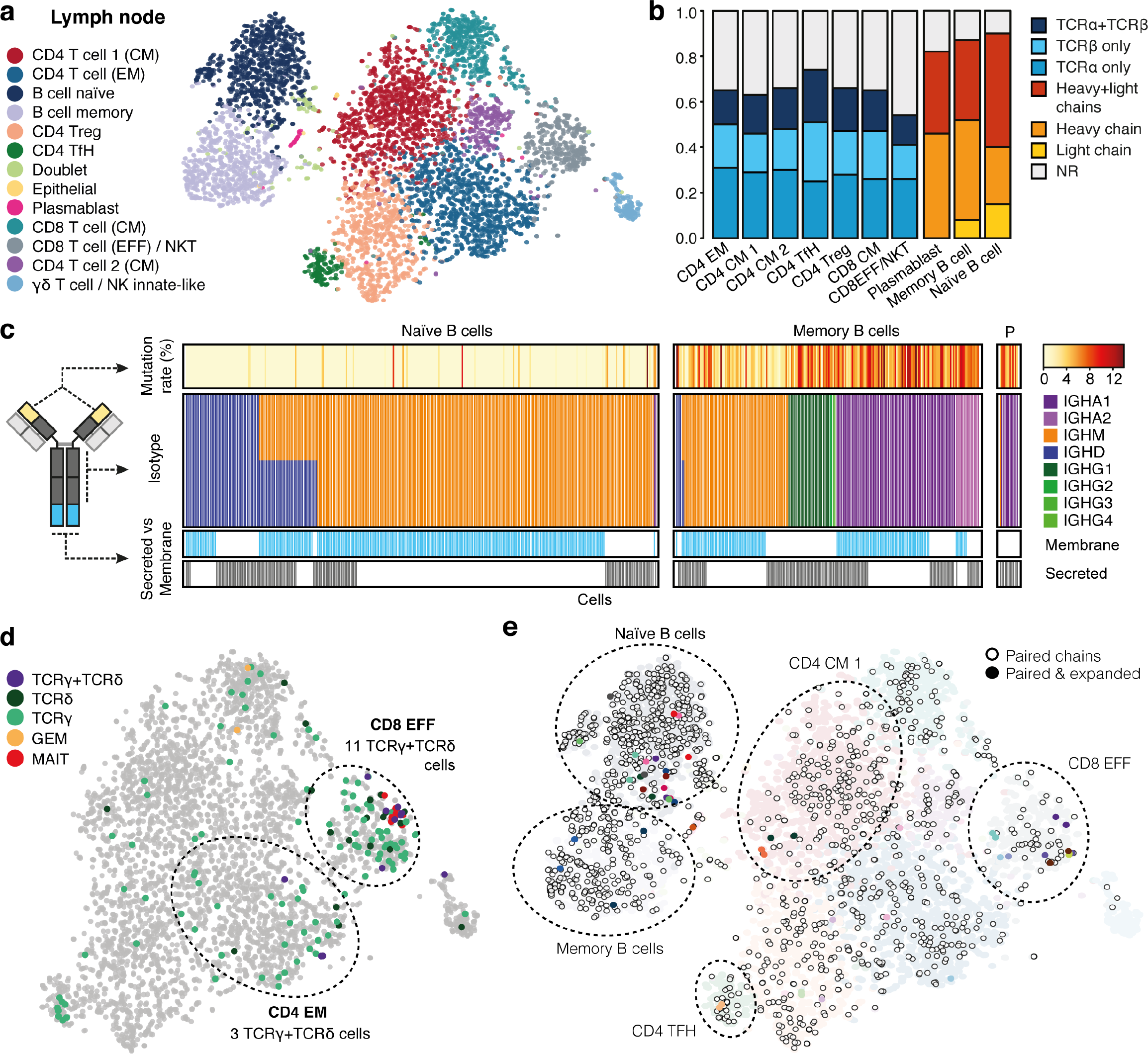
RAGE-Seq on a human lymph node. tSNE plot associated with 3’ gene expression profiling of 6,027 lymph node cells captured on the 10X Chromium platform. Number of cells: B cell memory, 738; B cell naïve, 853; CD4 EM, 1069; CD4 CM1, 1096; CD4 CM 2, 226; TfH, 142; Treg, 740; CD8 CM, 487; CD8EFF*/*NKT 405, 405; Plasmablast, 28; Innate-like, 144; Doublets, 86; Epithelial, 13. (**b**) Assignment of productive TCR and BCR clonotypes to each population identified in (a). (**c**) Characterisation of *IGH* mRNA contigs assigned to individual naïve (n=401) or Memory B cells (n=283) from the lymph node or Plasmablasts (P, n=15) from a matched tumour (Supplementary Fig. 6) Mutation rate (%) measures the percentage of the total number nucleotides in the V region mutated from germline. (**d**) Assignment of TCR*γ* and TCR *δ* clonotypes to the T cell compartments of the lymph node in (a). 92 T cells were assigned TCR*γ* clonotypes alone, 14 T cells were assigned TCR*6* clonotypes alone and 11 T cells were assigned paired TCR*γδ* chains. (**e**) Visualisation of the tSNE plot in (a) for cells assigned paired BCR (n=689) or paired TCR (n=705) chains and amongst these cells those clones that are expanded. Clones were considered expanded if a paired TCR or BCR sequence was found in more than one cell. 13 T cell expanded clones and 13 B cell expanded clones were identified. Each clone was represented by 2 cells. Gene correlation between cells with the same receptor sequence vs cells with different receptor sequences (P=2.55E-11 (T cells), P=2.10E-7 (B cells), Wilcoxon signed-rank test, see Supplementary Fig. 5).

Naïve B cells predominantly co-express *IGH* mRNAs with identical V(D)J sequences at their 5’ end but different constant region sequences at the 3’ end, produced by alternative mRNA splicing of the V(D)J exon to either *IGHM* or *IGHD* exons, encoding receptors with the same antigen specificity but IgM or IgD isotype that have different trafficking and subtly different signalling. In the breast cancer lymph node most cells classified as naïve by gene expression pattern had *IGHM* or both *IGHM* and *IGHD* mRNAs (**Fig 5C**). The fact that *IGHD* was not detected in many *IGHM*-bearing naïve cells is consistent with *IGHM* mRNA being 10-fold more abundant than *IGHD* mRNA {Yuan, 1984}. Upon activation, B cells acquire point mutations in their *V(D)J* exon through somatic hypermutation and undergo DNA rearrangements that replace the *IGHM* and *IGHD* constant region exons with *IGHG, IGHE* or *IGHA* constant region exons to produce IgG, IgE and IgA antibody isotypes {Alt, 1980; Chaudhuri, 2004}. As expected, memory B cells in the lymph node had more point mutations in the V(D)J segment of their mRNA than naïve B cells and two-thirds had undergone isotype switching to *IGHA* or *IGHG* (**Fig. 5C)**.

For each *IGH* transcript isotype, alternative splicing at the 3’ end generates either membrane-bound or secreted forms of immunoglobulin, with the latter gradually predominating as activated or memory B cells differentiate into antibody-secreting plasmablast cells. The presence of shared V(D)J sequences in both membrane and secretory *IGH* isoforms was detected in many single naïve and memory B cells (**Fig. 5C**), consistent with previous evidence from pooled naïve cells {Yuan, 1984}. As expected, only the secretory spliced form was detected in plasmablasts or plasma cells in the lymph node and in the tumour from the same patient (see below and **Supplementary Fig. 6**), and most were assigned *IGHA1* isotypes. This is consistent with plasmablasts and plasma cells having differentiated into high-rate antibody secreting cells, and with the dominant switching to IgA in plasma cells of normal and neoplastic breast tissue {Sieinski, 1980; Hsu, 1981}.

Our targeted capture panel included probes against *TRG* and *TRD* genes allowing for the detection of γδ T cells, a poorly-explored class of unconventional T cells of substantial interest to studies of infection and tumour immunology. A total of 11 T cells in the lymph node were assigned paired TCRγ and TCRδ chains, the majority of which clustered in the CD8 effector cluster. We also recovered 92 T cells with only the TCRγ chain and 14 T cells with only the TCRδ chain, again the majority clustering in the CD8 T cell effector population (**Fig. 5D**). T cells assigned TCRγ chains alone were found to frequently co-express TCRα and TCRβ chains, consistent with the timing of TCRγ rearrangement {Joachims, 2006}. In contrast, T cells assigned paired TCRγ and TCRδ chains were not co-assigned TCRα or TCRβ chains suggesting that they are committed γδ T cells. We explored the identification of other unconventional T cells that can recognise non-peptide antigens based on their invariant TCR usage such as Mucosal Associated Invariant T (MAIT) cells {Lepore, 2014; Reantragoon, 2013} and Germline-Encoded Mycolyl lipid-reactive (GEM) T cells {Van Rhijn, 2013}. Ten T cells were found to carry MAIT-associated TCRs which clustered closely together in the CD8 effector population, while two T cells with GEM-associated TCR chains were found in the CD4 effector memory cluster (**Fig. 5D).**

Evidence for clonal expansion in the lymph node was uncommon, with shared *V(D)J* sequences only detected in pairs of cells. For B cells there were 13 cell pairs with the same BCR sequence, the majority of which segregated in the naïve B cell cluster, while for T cells there were also 13 cell pairs with shared TCR sequence that also clustered by cell type **(Fig 5E).** Individual cells belonging to a clone mapped close to each other on the tSNE plot suggesting that cells with the same receptor are more transcriptionally similar to each other. Indeed, B cell and T cell clones with the same receptor sequence present more similar gene expression profiles than non-clonally expanded B cells (P=2.10E-07, paired Wilcoxon test) and T cells (P=2.55E-11) when comparing their Jaccard similarity coefficient for the 250 most abundant genes chosen from within each respective cell type cluster.

### Cross-tissue repertoire analysis

An important application of RAGE-Seq is the ability to track clonally related T or B cells across tissues, to gain systems-level insights into the evolution of immune responses. One such application is the analysis of lymphocytes in a tumour and its draining lymph node, the presumptive site of antigen presentation and source of tumour-infiltrating lymphocytes (TILs). In parallel with the lymph node analysis above, we performed RAGE-Seq on the patient’s resected primary breast tumour. From a total of 2,493 captured cells, 909 T cells and 215 B cells were identified (**Supplementary Fig. 6)**. A substantial number of receptor chains were found to be shared by lymphocytes within each tissue: 34 light chains amongst 157 B-cells; 11 TCRα chains, 7 TCRβ chains and 5 TCRγ chains amongst 134 T-cells (**Fig 6A**). Some chains showed significant tissue-specific enrichment, with one *IGL* sequence (V: *IGLV4-69*, J: *IGLJ3*, CDR3: QTWGTGFWV) expressed by 27 tumour-resident B cells and plasmablasts (16.9% of all light chains), but undetected in the lymph node.

**Figure 6.**
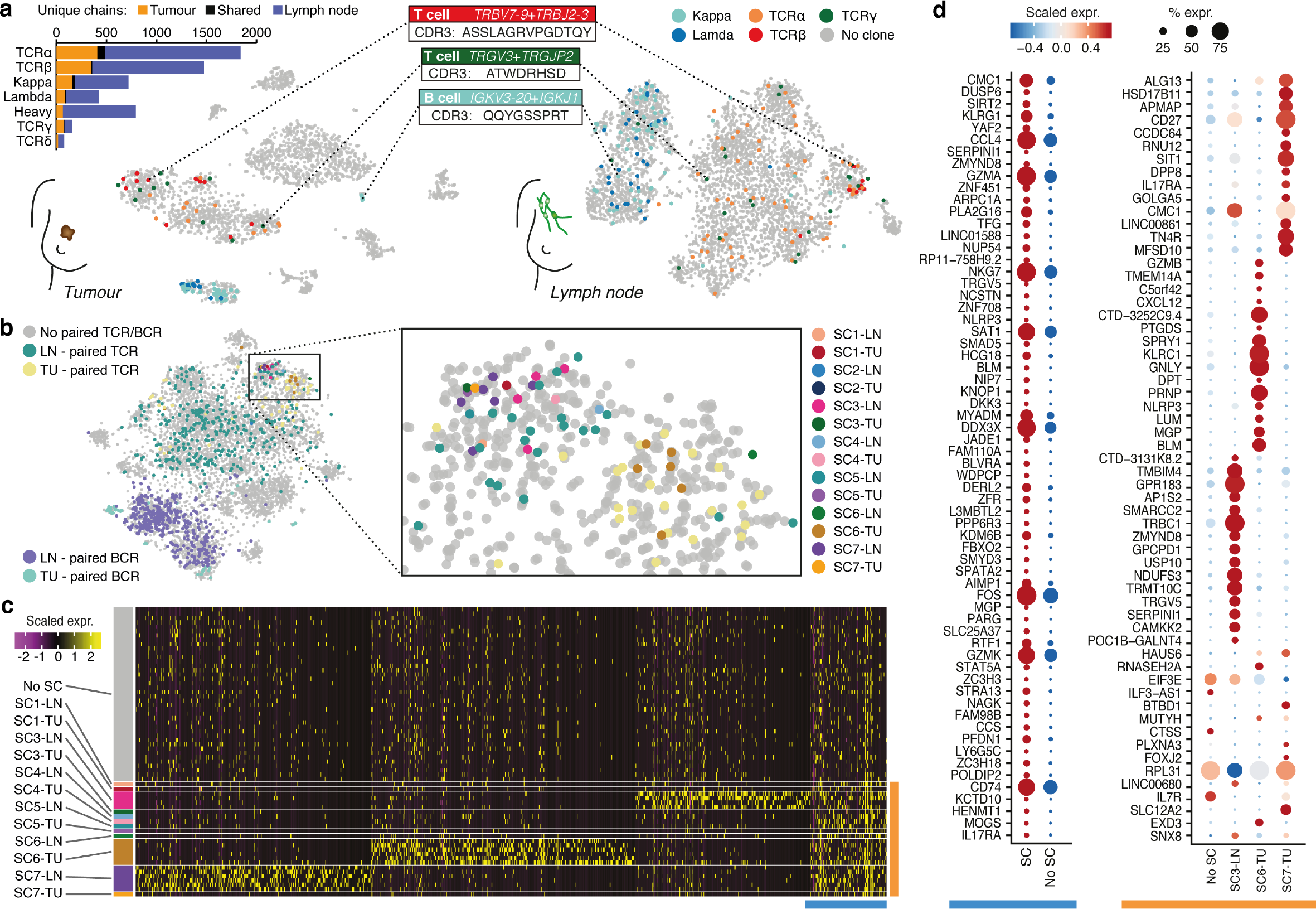
Repertoire analysis of patient matched lymph node and tumour. tSNE generated from 3’ 10x Chromium capture of patient matched tumour and lymph node. Select lymphocytes found to express shared chains across both tissues are highlighted. (**b**) tSNE plot of tumour and lymph node integrated using the canonical correlation algorithm of Seurat Satija et al. 2015. Analysis of TCR*β* and IGH mRNA contig sequences recovered on all lymphocytes shows 7 shared clonotypes between tumour and lymph (30 cells). Of those, a shared nearest neighbour (SNN) graph placed 6 together within the same cluster (CD8 T cell effector) irrespective of tissue origin. (**c**) Differentially expressed genes performed on all cells within the CD8 cluster. The non-shared cluster was randomly down sampled to 50 cells solely for visualisation purposes (heatmap in top panel). Of 1,328 differential expressed genes (P≤0.01, Wilcoxon signed-rank test), the top 65 were visualised using a dotplot (**d**) for cells with shared TCR sequences vs cells with unique sequences (left panel), and between groups of 3 or more cells expressing shared TCR sequences (right panel).

To investigate whether clonally related lymphocytes across the tumour and lymph node have common gene expression features, we analysed clones expressing shared paired receptor chains or single TCRβ or immunoglobulin heavy chains between three or more cells. We identified seven clones shared between the tumour and draining lymph node, six of them within the CD8 T cell effector cluster (**Fig 6B**). Differential gene expression analysis between cells belonging to a shared clone versus all others revealed a discrete gene signature for the clonally expanded cells compared to non-expanded clones, irrespective of tissue origin (**Fig 6C**). Cells belonging to shared clones expressed genes highly expressed in active tissue resident cytotoxic lymphocytes such *CCL4*, *NKG7*, *GZMA*, *GZMK* {Cheuk, 2017}, suggesting that they were activated by antigen exposure. Interestingly, we also observed sets of genes that were uniquely expressed by each of the clones (**Fig 6C**).

The presence of clonally expanded T cells between tissues suggested that these cells were proliferating in response to antigen stimulation. To examine this further, we used the scRNA-Seq data to perform cell cycle analysis of all cells within each CD8 T cell effector cluster of tumour and lymph node to infer whether TIL persistence of the clone is through proliferation occurring at the site of each sample, or through trafficking between tissues. A large proportion of T cells were in S, G2 or M phase, suggesting ongoing proliferation. Interestingly, proliferation of expanded clones in the tumour was found to be comparable to non-expanded clones. **(Supplementary Fig 6)**.

## Discussion

Single cell methods are needed to capture cell diversity arising at both ends of individual mRNA molecules, such as pairing the 5’ clonotype-specific V(D)J sequence of BCR or TCR transcripts with different 3’ sequences for secreted or membrane forms of different immunoglobulin isotypes, and with global expression profiling of 3’ ends. Here, we report a generalisable experimental workflow and computation pipeline to integrate gene expression with targeted analysis of splicing, structural variation and somatic mutation from thousands of single cells.

We have found RAGE-Seq to be robust in its ability to sample across both Illumina and Nanopore sequencing platforms and highly sensitive and accurate in providing full-length BCR and TCR sequences across immortalized and primary human B and T cells. Given its greater throughput and substantially lower cost, RAGE-Seq has significant advantages over SmartSeq2 for immune profiling. As a result, RAGE-Seq can circumvent the need to isolate specific lymphocyte populations by flow cytometry, permitting retrospective characterization of low abundance lymphocytes within tissues. We were able to identify clones with unique gene expression features that had expanded and were shared across tissues, despite unbiased sampling from a breast cancer, which generally have low TIL frequency. The capacity to screen large numbers of lymphocytes could have significant translational applications. Response to checkpoint inhibitors for immunotherapy has been linked to TCR clonality {Cha, 2014}, TIL frequency {Tumeh, 2014} and gene expression signatures {Ji, 2012}, yet these biomarkers have not been integrated at the single cell level. The application of RAGE-Seq to biopsies collected prior to and following treatment may accelerate the discovery of biomarkers or cell states that predict response to therapy. Additionally, the recovery of paired antigen receptors can be used with complimentary approaches to identify TCR ligands {Gee, 2018}, which could lead to novel neoantigen targets for chimeric antigen receptors to be built against {June, 2018}.

In this study we demonstrate the compatibility of RAGE-Seq with the 10X Chromium 3’ system, which should also be readily adaptable to any high-throughput single cell RNA-sequencing technologies that employ 3’ cell-barcode tagging {Macosko, 2015; Klein, 2015; Gierahn, 2017; Han, 2018; Fan, 2015; Rosenberg, 2018}. Recently, the commercially available Single Cell V(D)J + 5’ Gene Expression kit has been used to profile TILs in Breast cancer {Azizi, 2018} which relies on the incorporation of cell barcodes on the 5’ end of mRNA transcripts and V(D)J-specific PCR amplification. Compared to this method, RAGE-Seq has several advantages. It is compatible with DNA barcoded antibody technologies Abseq, CITE-Seq and REAP-Seq {Stoeckius, 2017; Peterson, 2017; Shahi, 2017}, which by allowing the additional measurement of cell surface proteins are powerful tools for immunophenotyping,. RAGE-Seq also sequences receptors from all lymphocytes in a single reaction, including γδ T cells which are of increasing interest in infection and cancer immunology {Zhao, 2018 #56}. RAGE-Seq provides full-length receptor sequence, which is greatly beneficial for the analysis of immunoglobulin somatic hypermutation and for the synthesis of recombinant antibodies, which can be used to explore the antigen specificity of B cells of interest.

A further advantage of RAGE-Seq is the ability to detect splice isoforms at the single cell level, which we have demonstrated by detecting *IGH* isoforms destined for antibody secretion or membrane-integration. To our knowledge, this is the first report integrating *IGH* V(D)J sequences with analysis of membrane or secreted exons. While we have relied on *de novo* assembly methods to generate splice isoform consensus sequences we believe that detection and quantification of isoforms will be enhanced with the future identification of UMI sequences from nanopore sequencing reads. A limitation of RAGE-Seq lies in the low recovery of cell barcodes (15-20%) due to the higher error-rate of base called nanopore sequencing data. We anticipate that bioinformatics tools leveraging raw nanopore signal {Wick, 2018} will increase this efficiency and reduce the number of long-reads that fail to be assigned to a cell.

While we have focused on lymphocyte receptors, any transcripts of interest can be targeted using variations of RAGE-Seq, simply by changing the composition of the capture probe library. One can envisage this method being applied to multiple areas of biology, such as oncology where variants of RAGE-Seq could be used to track the transcriptional consequences of clonal evolution at single cell resolution. The adaptability of RAGE-Seq across multiple scRNA-seq platforms and the flexibility to target a range of genes offers a new genomic toolkit that, amongst other applications, may be of particular value in comprehensively describing a human cell atlas {Regev, 2017}.

## Acknowledgements

We acknowledge the assistance of Chia-Ling Chan and Sunny Wu with single cell captures, David Koppstein with computational analysis, and Joseph Powell for stimulating discussions. This work was funded by The National Breast Cancer Foundation, John & Deborah McMurtrie, and The Kinghorn Foundation.

## References

Afik S, Yates KB, Bi K, Darko S, Godec J, Gerdemann U, et al. Targeted reconstruction of T cell receptor sequence from single cell RNA-seq links CDR3 length to T cell differentiation state. Nucleic acids research. 2017;45(16):e148.

Alt FW, Bothwell AL, Knapp M, Siden E, Mather E, Koshland M, et al. Synthesis of secreted and membrane-bound immunoglobulin mu heavy chains is directed by mRNAs that differ at their 3′ ends. Cell. 1980;20(2):293-301.

Azizi E, Carr AJ, Plitas G, Cornish AE, Konopacki C, Prabhakaran S, et al. Single-Cell Map of Diverse Immune Phenotypes in the Breast Tumor Microenvironment. Cell. 2018.

Bassing CH, Swat W, Alt FW. The mechanism and regulation of chromosomal V(D)J recombination. Cell. 2002;109 Suppl:S45-55.

Baillie JK, Barnett MW, Upton KR, Gerhardt DJ, Richmond TA, De Sapio F, et al. Somatic retrotransposition alters the genetic landscape of the human brain. Nature. 2011;479(7374):534-7.

Byrne A, Beaudin AE, Olsen HE, Jain M, Cole C, Palmer T, DuBois RM, Forsberg EC, Akeson M, Vollmers C. Nanopore long-read RNAseq reveals widespread transcriptional variation among the surface receptors of individual B cells. Nature Communications. 2017 Jul 19;8:16027.

Calis JJ, Rosenberg BR. Characterizing immune repertoires by high throughput sequencing: strategies and applications. Trends in immunology. 2014;35(12):581-90.

Camacho C, Coulouris G, Avagyan V, Ma N, Papadopoulos J, Bealer K, et al. BLAST+: architecture and applications. BMC bioinformatics. 2009;10:421.

Cao J, Packer JS, Ramani V, Cusanovich DA, Huynh C, Daza R, et al. Comprehensive single-cell transcriptional profiling of a multicellular organism. Science (New York, NY). 2017;357(6352):661-7.

Carlson CS, Emerson RO, Sherwood AM, Desmarais C, Chung MW, Parsons JM, et al. Using synthetic templates to design an unbiased multiplex PCR assay. Nature communications. 2013;4:2680.

Chaudhuri J, Alt FW. Class-switch recombination: interplay of transcription, DNA deamination and DNA repair. Nature reviews Immunology. 2004;4(7):541-52.

Chapman CJ, Zhou JX, Gregory C, Rickinson AB, Stevenson FK. VH and VL gene analysis in sporadic Burkitt’s lymphoma shows somatic hypermutation, intraclonal heterogeneity, and a role for antigen selection. Blood. 1996;88(9):3562-8.

Cheuk S, Schlums H, Sérézal IG, Martini E, Chiang SC, Marquardt N, Gibbs A, Detlofsson E, Introini A, Forkel M, Höög C. CD49a expression defines tissue-resident CD8+ T cells poised for cytotoxic function in human skin. Immunity. 2017 Feb 21;46(2):287-300.

Di Noia JM, Neuberger MS. Molecular mechanisms of antibody somatic hypermutation. Annual review of biochemistry. 2007;76:1-22.

Eltahla AA, Rizzetto S, Pirozyan MR, Betz-Stablein BD, Venturi V, Kedzierska K, Lloyd AR, Bull RA, Luciani F. Linking the T cell receptor to the single cell transcriptome in antigen-specific human T cells. Immunology and cell biology. 2016 Jul;94(6):604-11.

Fan HC, Fu GK, Fodor SP. Expression profiling. Combinatorial labeling of single cells for gene expression cytometry. Science (New York, NY). 2015;347(6222):1258367.

Gee MH, Han A, Lofgren SM, Beausang JF, Mendoza JL, Birnbaum ME, et al. Antigen Identification for Orphan T Cell Receptors Expressed on Tumor-Infiltrating Lymphocytes. Cell. 2018;172(3):549-63.e16.

Gierahn TM, Wadsworth MH, 2nd, Hughes TK, Bryson BD, Butler A, Satija R, et al. Seq-Well: portable, low-cost RNA sequencing of single cells at high throughput. Nature methods. 2017;14(4):395-8.

Han X, Wang R, Zhou Y, Fei L, Sun H, Lai S, et al. Mapping the Mouse Cell Atlas by Microwell-Seq. Cell. 2018;173(5):1307.

Hsu SM, Raine L, Nayak RN. Medullary carcinoma of breast: an immunohistochemical study of its lymphoid stroma. Cancer. 1981;48(6):1368-76.

Jain M, Koren S, Miga KH, Quick J, Rand AC, Sasani TA, et al. Nanopore sequencing and assembly of a human genome with ultra-long reads. Nature biotechnology. 2018;36(4):338-45.

Joachims ML, Chain JL, Hooker SW, Knott-Craig CJ, Thompson LF. Human alpha beta and gamma delta thymocyte development: TCR gene rearrangements, intracellular TCR beta expression, and gamma delta developmental potential–differences between men and mice. Journal of immunology (Baltimore, Md: 1950). 2006;176(3):1543-52.

June CH, Sadelain M. Chimeric antigen receptor therapy. New England Journal of Medicine. 2018 Jul 5;379(1):64-73.

Klein AM, Mazutis L, Akartuna I, Tallapragada N, Veres A, Li V, et al. Droplet barcoding for single-cell transcriptomics applied to embryonic stem cells. Cell. 2015;161(5):1187-201.

Laydon DJ, Bangham CR, Asquith B. Estimating T-cell repertoire diversity: limitations of classical estimators and a new approach. Philosophical transactions of the Royal Society of London Series B, Biological sciences. 2015;370(1675).

Lefranc MP, Giudicelli V, Duroux P, Jabado-Michaloud J, Folch G, Aouinti S, et al. IMGT(R), the international ImMunoGeneTics information system(R) 25 years on. Nucleic acids research. 2015;43(Database issue):D413-22.

Lepore M, Kalinichenko A, Colone A, Paleja B, Singhal A, Tschumi A, et al. Parallel T-cell cloning and deep sequencing of human MAIT cells reveal stable oligoclonal TCRbeta repertoire. Nature communications. 2014;5:3866.

Li H. Minimap2: pairwise alignment for nucleotide sequences. Bioinformatics (Oxford, England). 2018.

Li H, Handsaker B, Wysoker A, Fennell T, Ruan J, Homer N, et al. The Sequence Alignment/Map format and SAMtools. Bioinformatics (Oxford, England). 2009;25(16):2078-9.

Macosko EZ, Basu A, Satija R, Nemesh J, Shekhar K, Goldman M, et al. Highly Parallel Genome-wide Expression Profiling of Individual Cells Using Nanoliter Droplets. Cell. 2015;161(5):1202-14.

Mercer TR, Clark MB, Crawford J, Brunck ME, Gerhardt DJ, Taft RJ, et al. Targeted sequencing for gene discovery and quantification using RNA CaptureSeq. Nature protocols. 2014;9(5):989-1009.

Nestorowa S, Hamey FK, Sala BP, Diamanti E, Shepherd M, Laurenti E, Wilson NK, Kent DG, Göttgens B. A single cell resolution map of mouse haematopoietic stem and progenitor cell differentiation. Blood. 2016 Jan 1:blood-2016.

Peterson VM, Zhang KX, Kumar N, Wong J, Li L, Wilson DC, et al. Multiplexed quantification of proteins and transcripts in single cells. Nature biotechnology. 2017;35(10):936-9.

Picelli S, Bjorklund AK, Faridani OR, Sagasser S, Winberg G, Sandberg R. Smart-seq2 for sensitive full-length transcriptome profiling in single cells. Nature methods. 2013;10(11):1096-8.

Picelli S, Faridani OR, Bjorklund AK, Winberg G, Sagasser S, Sandberg R. Full-length RNA-seq from single cells using Smart-seq2. Nature protocols. 2014;9(1):171-81.

Potter KN, Hobby P, Klijn S, Stevenson FK, Sutton BJ. Evidence for involvement of a hydrophobic patch in framework region 1 of human V4-34-encoded Igs in recognition of the red blood cell I antigen. Journal of immunology (Baltimore, Md: 1950). 2002;169(7):3777-82.

Rizzetto S, Eltahla AA, Lin P, Bull R, Lloyd AR, Ho JWK, et al. Impact of sequencing depth and read length on single cell RNA sequencing data of T cells. Scientific reports. 2017;7(1):12781.

Rizzetto S, Koppstein DNP, Samir J, Singh M, Reed JH, Cai CH, et al. B-cell receptor reconstruction from single-cell RNA-seq with VDJPuzzle. Bioinformatics (Oxford, England). 2018.

Sale JE, Neuberger MS. TdT-accessible breaks are scattered over the immunoglobulin V domain in a constitutively hypermutating B cell line. Immunity. 1998;9(6):859-69.

Satija R, Farrell JA, Gennert D, Schier AF, Regev A. Spatial reconstruction of single-cell gene expression data. Nature biotechnology. 2015 May;33(5):495.

Shahi P, Kim SC, Haliburton JR, Gartner ZJ, Abate AR. Abseq: Ultrahigh-throughput single cell protein profiling with droplet microfluidic barcoding. Scientific reports. 2017;7:44447.

Shannon P, Markiel A, Ozier O, Baliga NS, Wang JT, Ramage D, Amin N, Schwikowski B, Ideker T. Cytoscape: a software environment for integrated models of biomolecular interaction networks. Genome research. 2003 Nov 1;13(11):2498-504.

Shugay M, Britanova OV, Merzlyak EM, Turchaninova MA, Mamedov IZ, Tuganbaev TR, et al. Towards error-free profiling of immune repertoires. Nature methods. 2014;11(6):653-5.

Sieinski W. Immunohistological patterns of immunoglobulins in dysplasias, benign neoplasms and carcinomas of the breast. Tumori. 1980;66(6):699-711.

Stoeckius M, Hafemeister C, Stephenson W, Houck-Loomis B, Chattopadhyay PK, Swerdlow H, et al. Simultaneous epitope and transcriptome measurement in single cells. Nature methods. 2017;14(9):865-8.

Stubbington MJT, Lonnberg T, Proserpio V, Clare S, Speak AO, Dougan G, et al. T cell fate and clonality inference from single-cell transcriptomes. Nature methods. 2016;13(4):329-32.

Reantragoon R, Corbett AJ, Sakala IG, Gherardin NA, Furness JB, Chen Z, et al. Antigen-loaded MR1 tetramers define T cell receptor heterogeneity in mucosal-associated invariant T cells. The Journal of experimental medicine. 2013;210(11):2305-20.

Regev A, Teichmann SA, Lander ES, Amit I, Benoist C, Birney E, et al. The Human Cell Atlas. Elife. 2017;6.

Rosenberg AB, Roco CM, Muscat RA, Kuchina A, Sample P, Yao Z, et al. Single-cell profiling of the developing mouse brain and spinal cord with split-pool barcoding. Science (New York, NY). 2018;360(6385):176-82.

Rudolph MG, Stanfield RL, Wilson IA. How TCRs bind MHCs, peptides, and coreceptors. Annual review of immunology. 2006;24:419-66.

Tirosh I, Izar B, Prakadan SM, Wadsworth MH, Treacy D, Trombetta JJ, Rotem A, Rodman C, Lian C, Murphy G, Fallahi-Sichani M. Dissecting the multicellular ecosystem of metastatic melanoma by single-cell RNA-seq. Science. 2016 Apr 8;352(6282):189-96.

Upadhyay AA, Kauffman RC, Wolabaugh AN, Cho A, Patel NB, Reiss SM, et al. BALDR: a computational pipeline for paired heavy and light chain immunoglobulin reconstruction in single-cell RNA-seq data. Genome medicine. 2018;10(1):20.

Van Rhijn I, Kasmar A, de Jong A, Gras S, Bhati M, Doorenspleet ME, et al. A conserved human T cell population targets mycobacterial antigens presented by CD1b. Nature immunology. 2013;14(7):706-13.

Wick RR, Judd, JM, Holt, KE. Deepbinner: Demultiplexing barcoded Oxford Nanopore reads with deep convolutional neural networks. bioRxiv. 2018 Jul 366526.

Xu JL, Davis MM. Diversity in the CDR3 region of V(H) is sufficient for most antibody specificities. Immunity. 2000;13(1):37-45.

Ye J, Ma N, Madden TL, Ostell JM. IgBLAST: an immunoglobulin variable domain sequence analysis tool. Nucleic acids research. 2013;41(Web Server issue):W34-40.

Yoshikai Y, Anatoniou D, Clark SP, Yanagi Y, Sangster R, Van den Elsen P, et al. Sequence and expression of transcripts of the human T-cell receptor beta-chain genes. Nature. 1984;312(5994):521-4.

Yuan D, Tucker PW. Regulation of IgM and IgD synthesis in B lymphocytes. I. Changes in biosynthesis of mRNA for mu- and delta-chains. Journal of immunology (Baltimore, Md: 1950). 1984;132(3):1561-5.

Zhang X, Chen MH, Wu X, Kodani A, Fan J, Doan R, et al. Cell-Type-Specific Alternative Splicing Governs Cell Fate in the Developing Cerebral Cortex. Cell. 2016;166(5):1147-62.e15.

Zheng GX, Terry JM, Belgrader P, Ryvkin P, Bent ZW, Wilson R, et al. Massively parallel digital transcriptional profiling of single cells. Nature communications. 2017;8:14049.

Zhao Y, Niu C, Cui J. Gamma-delta (gammadelta) T cells: friend or foe in cancer development? Journal of translational medicine. 2018;16(1):3.

Ziegenhain C, Vieth B, Parekh S, Reinius B, Guillaumet-Adkins A, Smets M, et al. Comparative Analysis of Single-Cell RNA Sequencing Methods. Molecular cell. 2017;65(4):631-43.e4.

